# Elevational filtering reshapes mountain understory communities through diversity decline, functional shifts, and species replacement

**DOI:** 10.64898/2026.11.04.764281

**Authors:** Syed Waseem Gillani, Mushtaq Ahmad, Muhammad Manzoor, Roberto Salguero-Gómez

## Abstract

**Aims:** Mountain understory communities are structured along elevational gradients, as environmental constraints influence which species and ecological strategies can persist. Yet, how community differentiation, local diversity, functional strategies, and species replacement vary along elevation remains poorly resolved. Here, we examine how elevation reorganizes herbaceous understory communities. We hypothesize that increasing elevation differentiates community composition, reduces α-diversity, favors persistence-oriented strategies, and drives β-diversity primarily through species turnover rather than nestedness.

**Location:** Kashmir Himalaya, Azad Jammu and Kashmir, Pakistan.

**Methods:** We examined 107 herbaceous species across 27 sites spanning 2,202– 3,696 m a.s.l. and three elevational zones: temperate, subalpine, and alpine. To test our hypotheses, we integrated analyses of community composition, α-diversity, functional composition, and β-diversity partitioning along the same elevational gradient.

**Results:** Our results reveal a coherent pattern of community reassembly with increasing elevation. Specifically, communities become increasingly distinct with elevational separation, while species richness and Shannon diversity decline toward higher elevations. Functional composition changes concurrently, with perennial and chamaephytic groups increasing in representation as annual and therophytic groups decline. By contrast, leaf-size responses are more restricted, with only leptophylls showing a significant positive elevational relationship. Notably, species turnover accounts for 97.95% of total Sørensen β-diversity and increases with elevational separation, whereas nestedness declines.

**Conclusions:** Our findings show that elevation reorganizes herbaceous understory communities through linked changes in local diversity, functional strategies, and species identity. Importantly, increasing community differentiation primarily reflects species replacement rather than progressive species loss. Together, these patterns are consistent with environmental filtering favoring persistence-oriented strategies, although the underlying mechanisms remain unresolved. Our study therefore contributes to understanding mountain understory community assembly by integrating taxonomic and functional differentiation along the same elevational gradient.

## 1. Introduction

Mountain landscapes provide natural laboratories to examine how environmental heterogeneity shapes plant community assembly (Stein et al. 2014; Perrigo et al. 2020). This terrain complexity creates substantial variation in ecological conditions across mountain landscapes for plant species (Stein et al. 2014). Moreover, fine-scale topographic variation can produce marked differences in thermal conditions, creating microhabitats for subalpine, alpine, and nival plants within short distances (Scherrer & Körner, 2011). As such, elevational gradients capture broader environmental variation across relatively short geographic distances, providing a useful setting for examining patterns of diversification and community organization (Perrigo et al. 2020; Testolin et al. 2021). Thus, understanding community variation along elevational gradients lays the foundation to identify the ecological processes underlying community assembly (Stein et al. 2014; Perrigo et al. 2020).

Environmental filtering can shape plant community composition and diversity along elevational gradients by sorting species according to their responses to changing environmental conditions (Bañares-de-Dios et al. 2024; Ma et al. 2024). Temperature and atmospheric conditions vary systematically with elevation (Körner, 2007), while local environmental factors, including soil properties, also change along mountain gradients (Hamid et al. 2021). Consequently, environmental filtering can sort species with different environmental responses from the regional pool into locally suitable communities (Zheng et al. 2022; Helm et al. 2024; López-Angulo et al. 2026). Such sorting can produce compositional differentiation among communities along elevation (Wani et al. 2023). Increasing environmental constraints can also limit local persistence and reduce α-diversity, the diversity of species within a single community (Whittaker, 1960), with declines in species richness and phylogenetic diversity reported along elevation (Galván-Cisneros et al. 2023). Yet, changes in taxonomic composition and diversity do not indicate whether environmental filtering also favors particular ecological strategies within communities (Klipel et al. 2023; Ratier Backes et al. 2023). Functional characteristics can therefore complement taxonomic responses by revealing whether the ecological strategies represented within communities vary along elevational gradients (Ferrara et al. 2024).

Life-history and vegetative characteristics provide complementary measures of plant ecological strategies associated with their persistence along elevational gradients (Junker & Larue-Kontić, 2018; Rosbakh & Poschlod, 2021). For instance, life span captures a temporal dimension of persistence (Chondol et al. 2023), whereas plant growth forms (Raunkiær, 1934) represent persistence strategies through the position and protection of renewal buds during unfavorable periods (Midolo et al. 2024). Beyond these temporal and bud-protection dimensions of persistence, vegetative characteristics capture additional variation in plant morphology associated with differences in elevational distributions (Binter et al. 2025). Thus, considering these dimensions jointly should reveal how elevational filtering reorganizes plant ecological strategies associated with persistence (Ferrara et al. 2024).

Plant communities can become compositionally differentiated along elevational gradients through species replacement or nested species loss (Samad et al. 2026). Such compositional variation among communities is captured by β-diversity (Anderson et al. 2011), which can be decomposed into turnover and nestedness (Baselga & Orme, 2012). Turnover refers to the replacement of species between assemblages, whereas nestedness reflects compositional differences in which species-poor assemblages form subsets of richer assemblages (*sensu* Baselga, 2010). Across ecosystems, turnover often contributes more to β-diversity than nestedness, although their relative contributions vary among ecological systems (Soininen et al. 2018). Along elevational gradients, environmental separation can promote turnover when communities differ in their environmental tolerances (Fontana et al. 2020). Partitioning β-diversity therefore allows elevational reassembly to be evaluated in terms of species replacement versus nested species loss (Wani et al. 2022).

Despite growing understanding of taxonomic, functional, and compositional responses to elevation, how these dimensions are linked within the same plant communities remains poorly resolved (Bañares-de-Dios et al. 2024). Evidence from multidimensional studies suggests that these facets of community structure do not necessarily respond in parallel, as changes in species composition may occur without corresponding changes in functional characteristics, and *vice versa* (Pescador et al. 2015; Klipel et al. 2023). Consequently, patterns observed in one dimension may not reliably indicate how other dimensions of community structure are being reorganized (Devictor et al. 2010; Cardoso et al. 2014). Resolving these relationships requires examining taxonomic, functional, and compositional responses together within the same elevational gradient (Schroeder et al. 2024). Such an integrated approach can reveal whether these responses vary independently or covary as linked components of community reassembly, providing a more complete understanding of how plant communities reorganize along elevation (Zheng et al. 2022).

Here, we assess how elevation shapes the composition, diversity, ecological strategies, and reassembly of herbaceous understory communities in the Himalayas. The Himalayas provide a suitable system to explore this question because its variation in elevation, aspect, and local habitat conditions is associated with marked spatial variation in plant diversity across the region (Hamid et al. 2021; Shaheen et al. 2023; Wani et al. 2023; Gillani et al. 2024). We test four hypotheses: (H1) Elevation progressively structures community composition, producing increasingly distinct assemblages from temperate through subalpine to alpine zones (Galván-Cisneros et al. 2023; Wani et al. 2023); (H2) α-diversity declines with elevation, resulting in lower species richness and abundance-weighted Shannon diversity toward alpine communities (Testolin et al. 2021); (H3) Communities shift toward persistence-oriented strategies with elevation, with greater perennial and protected or low-bud representation, lower annual, biennial, and therophytic representation, and a restructured community leaf-size spectrum (Chondol et al. 2023; Midolo et al. 2024); and (H4) Community reassembly occurs primarily through species turnover rather than nested species loss, with species replacement contributing more strongly to β-diversity and increasing with elevational separation (Fontana et al. 2020; Wani et al. 2023).

## 2. Methodology

To test our hypotheses on elevational community assembly, we examined herbaceous understory vegetation across 27 sites spanning temperate, subalpine, and alpine zones in the Himalayas (Figure 1A). We quantified community composition, α- and β-diversity along an elevational gradient between 2,202 and 3,696 m a.s.l., as well as functional structure. Briefly, we used multivariate community analyses to assess compositional differentiation and regression models to quantify diversity responses. We then analyzed abundance-weighted life span, Raunkiær growth form, and leaf-size composition to evaluate functional filtering. Finally, we partitioned Sørensen β-diversity into turnover and nestedness components to determine whether elevational reassembly primarily reflected species replacement or nested species loss.

**Figure 1.**
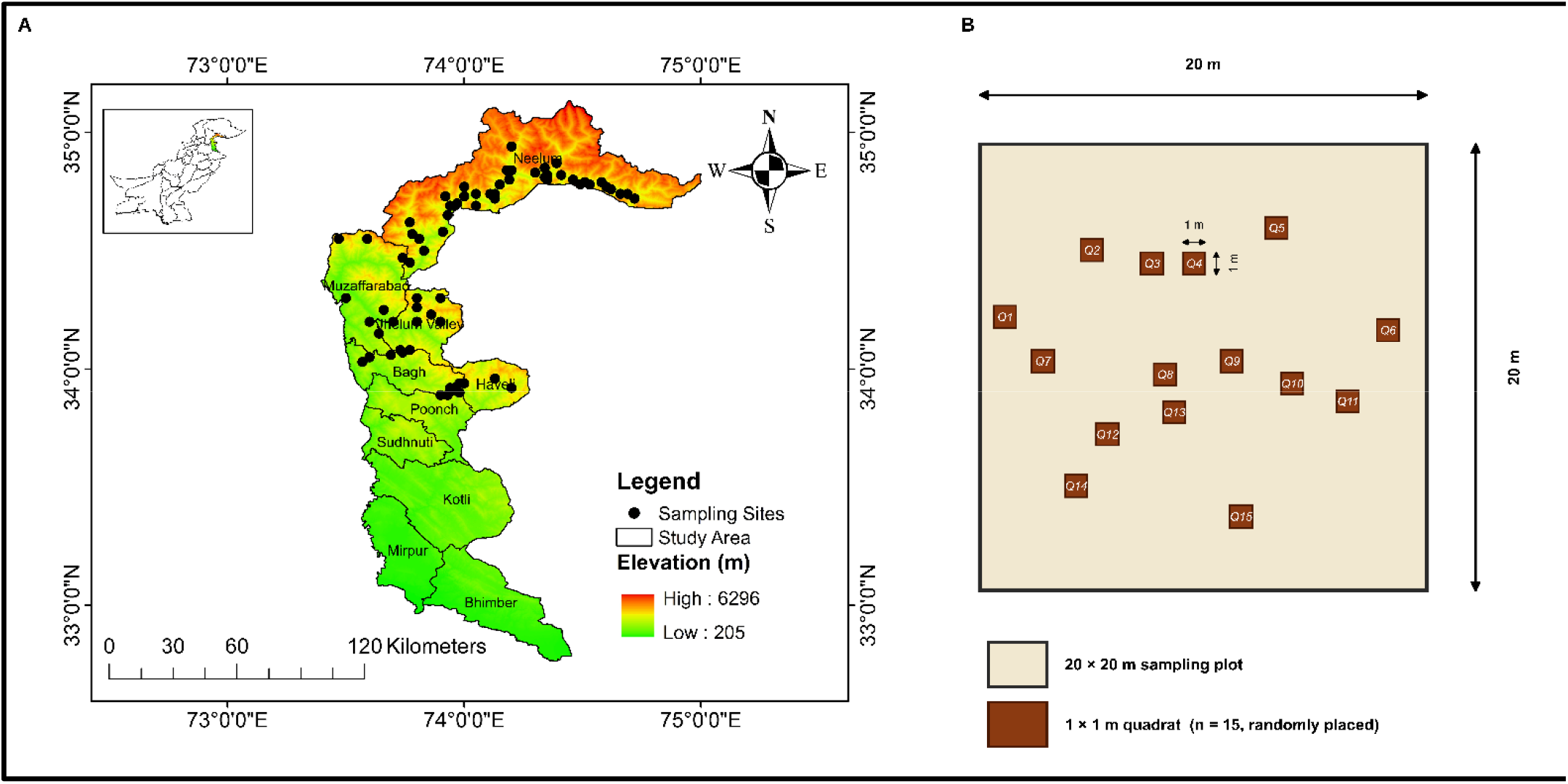
Study area and sampling design used to examine elevational variation in herbaceous understory communities in Kashmir Himalaya. **(A)** Geographical distribution of the 27 sampling sites along the elevational gradient, with black circles indicating sampling locations and the color gradient representing elevation. (B) Sampling design applied at each site, comprising a 20 × 20 m plot containing 15 randomly distributed 1 × 1 m quadrats for herbaceous understory vegetation sampling, providing a total sampled area of 15 m² per site.

### 2.1 Study area

To examine elevational variation in herbaceous understory communities, we conducted the study in Azad Jammu and Kashmir, Pakistan, within the western Himalayas (Figure 1A). Azad Jammu and Kashmir extends approximately from 33° to 36° N and 73° to 75° E and covers about 13,297 km² (GoAJK, 2019). The mountainous terrain in this region varies strongly in elevation, slope, aspect, and valley topography, creating pronounced environmental variation across the region (Khan et al. 2019). Elevation extends from low-lying southern valleys (360 m) to over 6,000 m in the northern mountains (GoAJK, 2014), with corresponding changes in vegetation structure and species composition (Khan et al. 2019; Shaheen et al. 2023).

Climate and vegetation vary markedly along this elevational gradient. Lower elevations experience a subtropical monsoon climate, whereas mid-elevations support moist temperate forests and higher elevations transition into subalpine and alpine vegetation (Shaheen et al. 2023). Across our sampling gradient, these vegetation transitions are represented by temperate, subalpine, and alpine communities. Temperatures decline with increasing elevation and frequently fall below freezing above 1,200 m during winter (Pak-Met, 2021). Annual precipitation averages approximately 1,300 mm, with much of the rainfall occurring during the July–September monsoon season (Pak-Met, 2021).

### 2.2 Vegetation sampling and data collection

To sample herbaceous understory vegetation along the elevational gradient, we selected 27 sites across temperate, subalpine, and alpine habitats of Azad Jammu and Kashmir following preliminary reconnaissance surveys in 2022. The selected sites were distributed across Neelum Valley, Jhelum Valley, Muzaffarabad, Haveli, and Bagh (Figure 1A). We then surveyed each site once during the main growing season (June– September), with fieldwork concentrated in 2023, when herbaceous vegetation was at peak vegetative growth, facilitating reliable species identification and vegetation assessment. We selected sites supporting herbaceous understory vegetation while representing variation in elevation, habitat, and local environmental conditions.

To standardize sampling effort across sites, we established a 20 × 20 m^2^ plot supporting herbaceous understory vegetation at each site. We randomly distributed fifteen 1 × 1 m^2^ quadrats within each plot (total sampled area of 15 m² per site; Figure 1 B). We recorded the identity and abundance of all herbaceous species within each quadrat and applied the same sampling design across the 27 sites. We identified species using the Flora of Pakistan (https://legacy.tropicos.org/Project/Pakistan) and regional floristic literature (Ali & Qaiser, 1986). We then verified accepted taxonomic names and synonyms using Plants of the World Online (https://powo.science.kew.org/), and the Royal Botanic Kew Gardens (https://www.kew.org/). Finally, to characterize topographic variation along the elevational gradient, we recorded elevation, latitude, longitude, slope, and aspect at each site. These variables provided the spatial and environmental context for testing hypotheses H1–H4 on herbaceous plant diversity and community composition. We measured geographic coordinates and elevation using a handheld GPS receiver and slope and aspect using a clinometer.

### 2.3 Community composition

To test our hypothesis H1, which predicts that elevation progressively structures community composition and produces increasingly distinct assemblages from temperate through subalpine to alpine zones, we quantified compositional variation across the 27 study sites. To do so, we calculated pairwise Bray–Curtis dissimilarities from the recorded species abundances. Bray–Curtis dissimilarity quantifies abundance-based differences in community composition using non-negative species data (Bray & Curtis, 1957). We retained species abundances in their recorded form because the measurements comprised comparable abundance or cover values across sites. The resulting Bray–Curtis dissimilarity matrix provided the basis for our multivariate analyses.

To assess compositional differentiation among sites along the elevational gradient, we used two-dimensional non-metric multidimensional scaling (NMDS). We fitted the ordination to Bray–Curtis dissimilarities (Bray & Curtis, 1957) using multiple random starts and assessed ordination adequacy based on the final stress value (Kruskal, 1964). We then tested whether continuous elevation explained variation in community composition using permutational multivariate analysis of variance (PERMANOVA) with 9,999 permutations (Anderson, 2001). Specifically, we fitted a separate PERMANOVA with elevational zone as a categorical predictor to assess compositional differentiation among temperate, subalpine, and alpine communities. Because differences in within-group dispersion can influence PERMANOVA (Anderson & Walsh, 2013), we tested homogeneity of multivariate dispersion among zones using PERMDISP with 9,999 permutations before interpreting the zone effect (Anderson, 2006).

To test the directional prediction of H1 that community dissimilarity increases with elevational separation, we related Bray–Curtis dissimilarity to absolute pairwise elevational distance. To do so, we used a Mantel test with 9,999 permutations (Mantel, 1967). We based statistical inference on permutation testing because our 351 pairwise site comparisons are not statistically independent. Next, we fitted a linear relationship to quantify the direction and magnitude of the distance-decay pattern. For visualization, we estimated uncertainty around the fitted relationship using 1,000 site-level bootstrap resamples. We conducted NMDS, PERMANOVA, PERMDISP, and Mantel analyses in R using the *vegan* package (Oksanen et al. 2026).

### 2.4 Species richness and diversity

To test our hypothesis H2, that increasing elevational constraints reduce local α-diversity, resulting in lower species richness and abundance-weighted Shannon diversity toward alpine communities, we quantified both diversity components across our 27 study sites. Specifically, we calculated species richness as the number of species with non-zero abundance at each site. We calculated the Shannon diversity index as *H*′ = −Σ *p* ln(*p*), where *p* denotes the relative abundance of species *i* at a site and ln denotes the natural logarithm (Shannon, 1948).

We first modelled species richness using a Poisson generalized linear model (GLM) with a log link. We assessed overdispersion using the Pearson chi-square statistic divided by the residual degrees of freedom and considered a negative-binomial GLM if the statistic indicated substantial overdispersion. However, the dispersion statistics were below 1, indicating no evidence of overdispersion (Table S3). We therefore retained the Poisson GLM for the final analysis rather than fitting a negative-binomial model. For the Shannon diversity index, we initially fitted an ordinary least-squares model and evaluated its residual assumptions. Specifically, we assessed residual normality using the Shapiro–Wilk test and homoscedasticity using the Breusch– Pagan test. Because the residuals violated the assumption of homoscedasticity (Figure S1), we subsequently fitted a Gamma GLM with a log link as the primary model for Shannon diversity.

Finally, we tested whether both diversity metrics showed predominantly monotonic or nonlinear responses to elevation. Although H2 predicts a monotonic decline, plant species richness in mountains can either decline with elevation or peak at intermediate elevations (Rahbek, 1995; Grytnes & Vetaas, 2002; Rahbek, 2005). A mid-elevation peak would therefore contradict the progressive decline predicted by H2. To assess these alternative responses, we compared linear models with corresponding models containing a quadratic elevation term, using the same error distribution and log link (Poisson for species richness; Gamma for Shannon diversity). We retained the quadratic model only when it reduced the Akaike Information Criterion (AIC) by more than two units relative to the linear model and a nested-model test of the quadratic term was statistically significant (likelihood-ratio test for richness; *F* test for Shannon diversity; *P* < 0.05) (Burnham & Anderson, 2002; Table S4). Following Nakagawa & Cuthill (2007), we quantified elevational effects using model coefficients, standard errors, 95% Wald confidence intervals, and two-sided *P*-values. To illustrate the fitted relationships, we generated model predictions with associated 95% confidence intervals across the observed elevational range (2,202–3,696 m).

To distinguish the elevational communities considered in H1, we classified the 27 study sites into temperate, subalpine, and alpine zones. We used these zones to distinguish sites by color in the visualizations (Figs. 1, 3A–B, and S1). The zones represent non-overlapping elevation classes (temperate, 2,202–2,925 m; subalpine, 3,144–3,417 m; alpine, 3,526–3,696 m), containing 17, six, and four sites, respectively. However, we excluded elevational zone as a predictor from the primary diversity models testing H2 for two reasons. First, zone membership explains 80.5% of the variance in elevation, indicating a strong association between the two predictors that could introduce collinearity if included together (Dormann et al. 2013). Second, categorizing elevation would discard within-zone variation and could reduce statistical power (Royston et al. 2006). Accordingly, we retained elevation as a continuous predictor to preserve information from all 27 sites and directly test the progressive decline predicted by H2. To assess the robustness of these relationships, we also fitted models with elevational zone as a covariate. The elevational slopes remained negative and statistically significant for both diversity metrics (Table S5). We conducted all analyses in R (R Core Team, 2026), using *stats* for model fitting and *lmtest* for model diagnostics (Zeileis & Hothorn, 2002).

### 2.5 Functional composition

To test our hypothesis H3, that increasing elevation filters understory communities toward persistence-oriented ecological strategies, we quantified abundance-weighted functional composition across 27 study sites. We predicted increasing representation of perennial species and Raunkiær growth form with protected or low-positioned perennating buds (chamaephytes, hemicryptophytes, and geophytes), alongside declining representation of annual and biennial species and therophytes (Rosbakh & Poschlod, 2021; Midolo et al. 2024). We also predicted a shift toward smaller leaf-size classes with increasing elevation, consistent with their reduced susceptibility to frost damage in cold environments (Wright et al. 2017). To quantify these responses, we converted species abundances to site-level proportions for three trait dimensions: life span (annual, biennial, and perennial), Raunkiær growth form (therophyte, chamaephyte, hemicryptophyte, and geophyte), and leaf-size class (leptophyll, nanophyll, microphyll, mesophyll, and macrophyll). Given that proportions within each trait dimension sum to one, we treated functional composition as compositional data rather than as independent response variables (Aitchison, 1986).

Raunkiær growth form was defined by the position of perennating buds during unfavorable periods. Therophytes survive unfavorable periods as seeds; chamaephytes retain buds close to the ground (<25 cm); hemicryptophytes retain buds at or near the soil surface; and geophytes maintain perennating organs below ground (Raunkiær, 1934; Midolo et al. 2024). Leaf-size classes followed Raunkiær (1934), as summarized by Baladrón et al. (2023), and were defined by leaf area: leptophyll (<25 mm²), nanophyll (25–225 mm²), microphyll (225–2,025 mm²), mesophyll (2,025–18,225 mm²), and macrophyll (18,225–164,025 mm²).

To prepare the trait proportions for subsequent log-ratio analysis, which requires strictly positive values, we first replaced zeros and then transformed the compositional data for multivariate analysis. Specifically, within each trait dimension, we treated zeros as proportions too small to be quantified in our samples and replaced them with small positive values derived from the observed non-zero values, using multiplicative replacement, which multiplies all non-zero components by a common factor and thereby preserves the ratios among them (Martín-Fernández et al. 2003). We then closed each composition by rescaling its components so that they again summed to one. We transformed the resulting compositions using the centred log-ratio (clr) transformation (Aitchison, 1986). As clr coordinates sum to zero and are therefore linearly dependent, we subsequently represented each trait dimension using independent isometric log-ratio (ilr) coordinates for our primary multivariate analyses (Egozcue et al. 2003; Pawlowsky-Glahn et al. 2015).

We next tested whether functional composition changes with continuous elevation and differs among elevational zones. To do so, we fitted separate multivariate analyses of variance (MANOVAs) to the ilr coordinates of each trait dimension and evaluated multivariate effects using Pillai’s trace. We analyzed continuous elevation and elevational zone in separate models because the three zones represent categorical classifications of the same elevational gradient. To control the false discovery rate across the three trait dimensions, we adjusted multivariate *P*-values using the Benjamini–Hochberg procedure (Benjamini & Hochberg, 1995).

Finally, we identified the individual trait classes underlying the multivariate responses. We modelled clr-transformed proportions of each trait class against elevation using linear models. We assessed differences among elevational zones using separate linear models followed by estimated marginal mean comparisons where appropriate. We adjusted the resulting *P*-values within each trait dimension using the Benjamini–Hochberg procedure. For trait classes containing zeros at ≥30% of sites, we additionally used a two-part analysis to separate elevational changes in occurrence probability from changes in relative abundance among occupied sites. For visualization, we generated elevational response curves from the fitted clr-based models, back-transformed them to relative trait proportions, and estimated uncertainty using 1,000 site-level bootstrap resamples. We implemented estimated marginal means and multiple comparisons in R using *emmeans* and *multcomp* and conducted bootstrap resampling using *boot* (Hothorn et al. 2008; Canty & Ripley, 2025).

### 2.6 β-diversity partitioning

To test hypothesis H4, whereby community differentiation should occurs primarily through species turnover rather than nested species loss, and that species replacement increases with elevational separation, we partitioned β-diversity across the 27 study sites. For this analysis, we converted species abundance data into a binary site × species incidence matrix to quantify compositional differences based on species occurrences rather than abundances. Following Baselga (2010), we partitioned total Sørensen β-diversity (βSOR) into turnover, measured as Simpson dissimilarity (βSIM), and nestedness-resultant dissimilarity (βSNE), such that βSOR = βSIM + βSNE. Values of βSOR range from zero, indicating identical species composition, to one, indicating no shared species. Within this partitioning framework, high βSIM indicates that compositional differences arise primarily from species replacement. In contrast, high βSNE indicates that differences are primarily attributable to nestedness, with the species-poorer assemblage tending to form a subset of the species-richer assemblage.

Accordingly, under H4, we expected turnover (βSIM) to contribute more than nestedness-resultant dissimilarity (βSNE) to overall β-diversity. We further predicted that pairwise βSIM would increase with elevational separation between sites. We calculated both multiple-site and pairwise β-diversity components using the Sørensen framework implemented in the betapart package (Baselga & Orme, 2012).

To examine how β-diversity components varied across the elevational gradient, we first quantified the relative contributions of turnover and nestedness to total β-diversity across all sites. We then calculated βSOR, βSIM, and βSNE for each of the 351 unique site pairs to characterize pairwise variation in β-diversity components. For each site pair, we quantified elevational separation as the absolute difference in elevation between the two sites. Finally, we related pairwise turnover and nestedness separately to elevational separation to assess whether increasing elevational separation corresponds to greater species replacement or nested species loss.

Finally, we accounted for the non-independence of pairwise comparisons when evaluating relationships with elevational separation. Because the 351 pairwise observations share sites, they do not represent independent sampling units, so we assessed statistical significance using site-label permutation tests with 9,999 permutations rather than conventional ordinary least-squares inference (Legendre & Legendre, 2012; Martínez-Santalla et al. 2022). We quantified the direction and magnitude of each relationship using a Mantel-type correlation coefficient (Mantel, 1967) and fitted a linear relationship to estimate the change in βSIM and βSNE per 1,000 m of elevational separation. For visualization, we estimated 95% confidence intervals around the fitted relationships using 1,000 site-level bootstrap resamples, which retained sites, rather than individual pairwise comparisons, as the sampling units (Martín-Devasa et al. 2022). We implemented the Sørensen β-diversity partitioning using the *betapart* R package (Baselga & Orme, 2012).

### 2.7 Statistical analysis and software

To ensure a reproducible analytical framework, we conducted all statistical analyses in R (R Core Team, 2026), with the 27 sampling sites representing the independent observational units. We used *vegan* for multivariate community analyses, including ordination, dissimilarity, and permutation-based analyses (Oksanen et al. 2026), and *betapart* for partitioning Sørensen β-diversity into turnover and nestedness components (Baselga & Orme, 2012). We implemented generalized linear models and related regression analyses using the base R *stats* package, with *MASS* for negative-binomial modelling where required (Venables & Ripley, 2002) and *lmtest* for regression diagnostics, including the Breusch–Pagan test (Breusch & Pagan, 1979; Zeileis & Hothorn, 2002). We used *emmeans* to estimate model-based marginal means and conduct associated comparisons (Lenth & Piaskowski, 2026), and *multcomp* for simultaneous multiple-comparison procedures (Hothorn et al. 2008). To quantify uncertainty while retaining sites as the independent sampling units, we implemented site-level bootstrap resampling using *boot* (Davison & Hinkley, 1997; Canty & Ripley, 2025). We imported data from Excel files using *readxl* (Wickham & Bryan, 2026), produced graphical outputs using *ggplot2* (Wickham, 2016), assembled multi-panel figures using *patchwork* (Pedersen, 2025), and applied additional graphical scaling and formatting using *scales* (Wickham et al. 2025).

## 3 Results

### 3.1 Elevational structuring of community composition

We found that elevation strongly structures understory plant community composition across the 27 study sites (Figure 2A–B; Table 1). Consistent with our hypothesis H1, temperate, subalpine, and alpine communities show clear compositional differentiation along the elevational gradient. Specifically, our non-metric multidimensional scaling (NMDS) ordination adequately represents the Bray–Curtis dissimilarity structure (stress = 0.109) and reveals separation among communities (Figure 2A). Similarly, elevation explains 26.8% of compositional variation (PERMANOVA, pseudo-*F*_₁, ₂₅_ = 9.133, *R*² = 0.268, *P* < 0.001), while elevational zones explain 36.5% (pseudo-*F*_₂, ₂₄_ = 6.894, *R*² = 0.365, *P* < 0.001; Figure 2A; Table 1).

**Figure 2.**
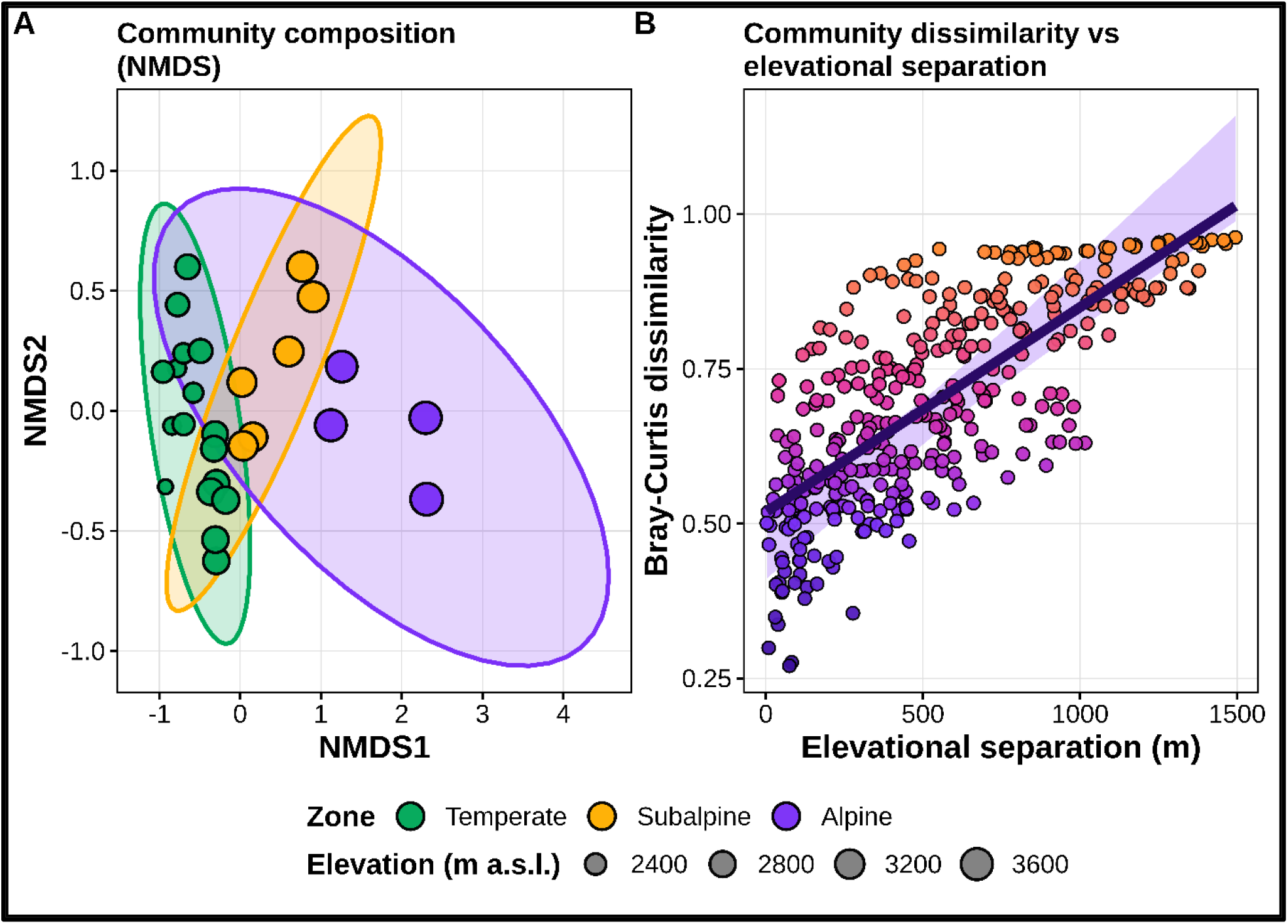
Understory plant communities become increasingly different as elevation increases and elevational distance increases among sites. **(A)** Non-metric multidimensional scaling (NMDS) ordination based on Bray–Curtis dissimilarity shows differences in community composition among temperate, subalpine and alpine sites. Sites plotted closer together contain more similar understory communities, whereas sites farther apart are more compositionally distinct. Colours identify the three elevational zones, dot size represents site elevation (m a.s.l.), and ellipses show the distribution of sites within each zone. The two-dimensional ordination has a stress value of 0.109. (**B**) Bray–Curtis dissimilarity increases with pairwise elevational separation, showing that sites farther apart in elevation support more compositionally distinct understory communities (Mantel *r* = 0.745, *P* < 0.001). Each point represents a pairwise comparison between two sites, the solid line shows the fitted relationship between elevational separation and community dissimilarity, and the shaded band shows the associated uncertainty.

**Table 1.** Elevation explains substantial variation in understory plant community composition across the study gradient. Separate permutational multivariate analysis of variance (PERMANOVA) models based on Bray–Curtis dissimilarity quantify the effects of elevation as a continuous gradient and elevational zone as a categorical predictor across 27 sites spanning 2,202–3,696 m a.s.l. Elevational zones comprise temperate (n = 17), subalpine (n = 6) and alpine (n = 4) sites. Both models use 9,999 permutations. *R*² represents the proportion of total variation in community composition explained by each predictor, pseudo-*F* represents the strength of compositional differentiation relative to residual variation, df denotes degrees of freedom, and the *P*-value indicates statistical significance.

| **Model** | **Predictor** | **df** | **Sum of squares** | ***R*<sup>2</sup>** | **Pseudo-*F*** | ***P*-value** | **Permutations** |
|---|---|---|---|---|---|---|---|
| Community composition | Elevation (continuous) | 1 | 1.788 | 0.268 | 9.133 | <0.001 | 9,999 |
| Community composition | Elevational zone | 2 | 2.439 | 0.365 | 6.894 | <0.001 | 9,999 |

Our analyses further show no significant differences in within-zone dispersion (PERMDISP, *F*_₂, ₂₄_ = 0.259, *P* = 0.769; Table S1). Moreover, community dissimilarity increases strongly with pairwise elevational separation (Mantel *r* = 0.745, *P* < 0.001; Figure 2B; Table S2). The fitted descriptive relationship indicates an increase of approximately 0.331 Bray–Curtis dissimilarity units per 1,000 m of elevational separation. Thus, our continuous and categorical analyses support H1, with greater elevational separation associated with increasingly distinct understory plant communities.

### 3.2 Elevational declines in α-diversity

Species richness declines significantly with increasing elevation across the 27 study sites (Figure 3A; Table 2). Consistent with H2, understory communities contain fewer species toward higher elevations. Specifically, species richness shows a negative elevational relationship (β = −0.268 per 1,000 m, SE = 0.075, 95% Wald CI = −0.414 to −0.121, *P* < 0.001; Figure 3A; Table 2). The Poisson model shows no overdispersion (Pearson χ²/df = 0.645; Table S3), while adding a quadratic term does not improve model fit (likelihood-ratio test, *P* = 0.730; ΔAIC = 1.880; Table S4). Thus, species richness declines with elevation, with no evidence of improved model fit from a quadratic term.

**Table 2.** Understory species richness and Shannon diversity decline significantly with increasing elevation. Separate generalized linear models (GLMs) quantify elevational effects on species richness and Shannon diversity across 27 study sites. Species richness is modelled using a Poisson GLM with a log link, whereas Shannon diversity is modelled using a Gamma GLM with a log link. Elevation coefficients (β) are expressed per 1,000 m increase in elevation, with negative values indicating declines toward higher elevations. SE denotes standard error, 95% CI denotes the 95% confidence interval, and *P*-values indicate statistical significance.

| **Response** | **Primary model** | **Elevation effect per 1,000 m (*β*)** | **SE** | **95% CI** | ***P*-value** |
|---|---|---|---|---|---|
| Species richness (S) | Poisson GLM (log link) | −0.268 | 0.075 | −0.414 to −0.121 | <0.001 |
| Shannon diversity (*H*′) | Gamma GLM (log link) | −0.089 | 0.028 | −0.143 to −0.035 | 0.003 |

We also found that Shannon diversity declines significantly with elevation (Figure 3B; Table 2). Specifically, the Gamma model reveals a negative relationship (β = −0.089 per 1,000 m, SE = 0.028, 95% Wald CI = −0.143 to −0.035, *P* = 0.003). Although the initial ordinary least-squares model shows no significant departure from residual normality (Shapiro–Wilk *P* = 0.744) but reveals significant heteroscedasticity (Breusch– Pagan *P* = 0.031; Figure S1). Consequently, we retain the Gamma model for the primary analysis (Table S3). As with species richness, the quadratic elevation term does not improve model fit (*F* test, *P* = 0.643; ΔAIC = 1.760; Table S4). Furthermore, both negative elevational relationships remain significant after accounting for zone (richness: β = −0.407, *P* = 0.012; Shannon diversity: β = −0.223, *P* < 0.001; Table S5). Thus, our results support H2 across the elevational gradient.

### 3.3 Elevational shifts in functional composition

Our results show significant shifts in functional composition along elevation and among temperate, subalpine, and alpine communities (Figure 4A–C; Table S6). Consistent with H3, communities shift toward persistence-oriented strategies, particularly in life span and life form. Specifically, multivariate analyses based on isometric log-ratio coordinates reveal significant elevational relationships for life span (Pillai’s trace = 0.549, *F* = 14.620, *P*adj < 0.001), life form (Pillai’s trace = 0.651, *F* = 14.315, *P*adj < 0.001), and leaf size (Pillai’s trace = 0.534, *F* = 6.297, *P*adj = 0.002). All three dimensions also differ among elevational zones (Table S5).

**Figure 3.**
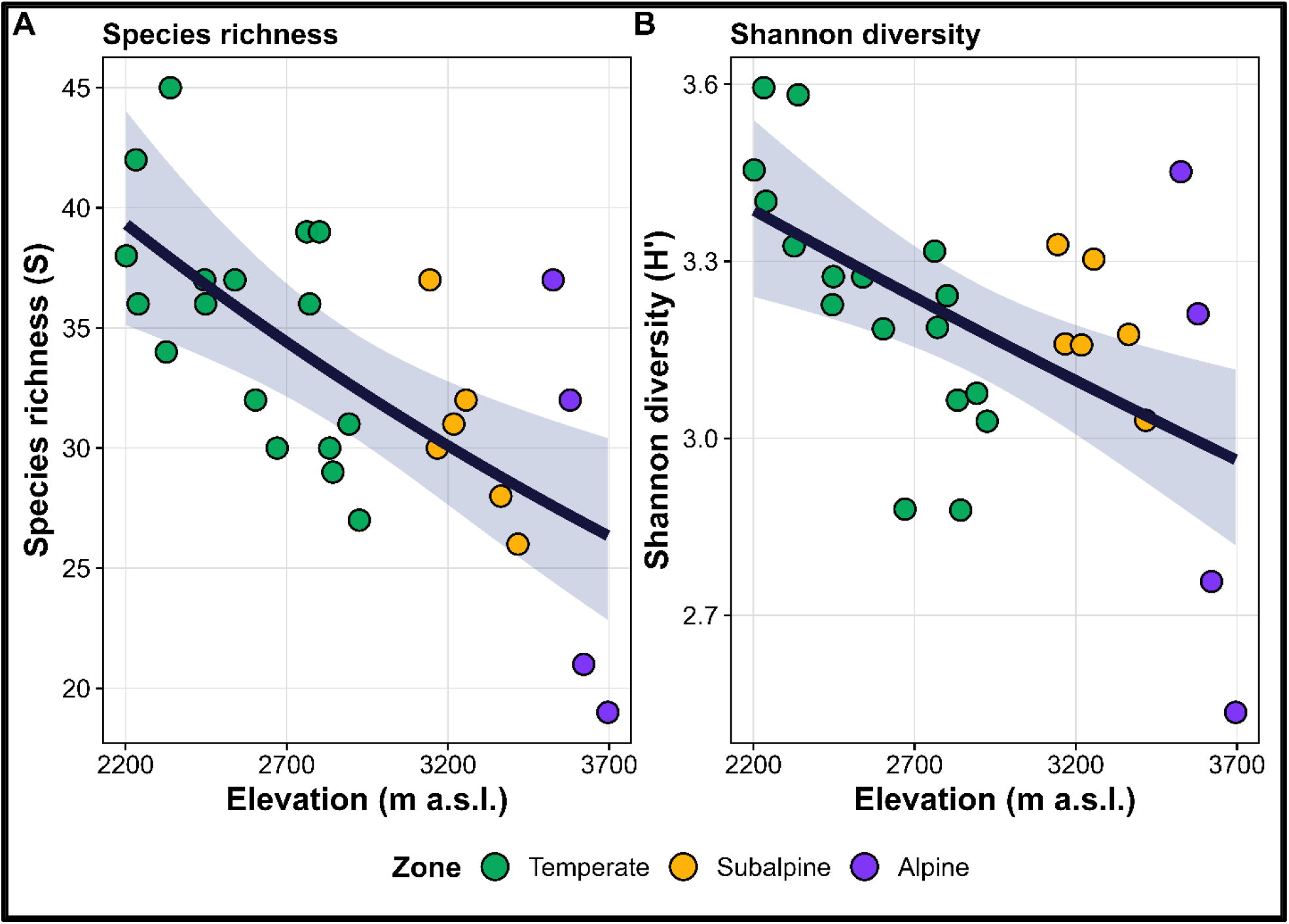
Understory species richness and Shannon diversity decline with increasing elevation. (**A**) Species richness, the number of species recorded at each site, decreases across the elevational gradient. The solid line shows the response predicted by the selected Poisson generalized linear model (GLM) with a log link. (**B**) Shannon diversity (*H*′), calculated from relative species abundances using the natural logarithm, also decreases with increasing elevation. The solid line shows the response predicted by the selected Gamma GLM with a log link. In both panels, dots represent the 27 study sites, with colours identifying temperate, subalpine and alpine zones. Elevational zone is shown for visual reference and is not included as a predictor in either model. Shaded bands represent 95% confidence intervals around the model-predicted relationships.

**Figure 4.**
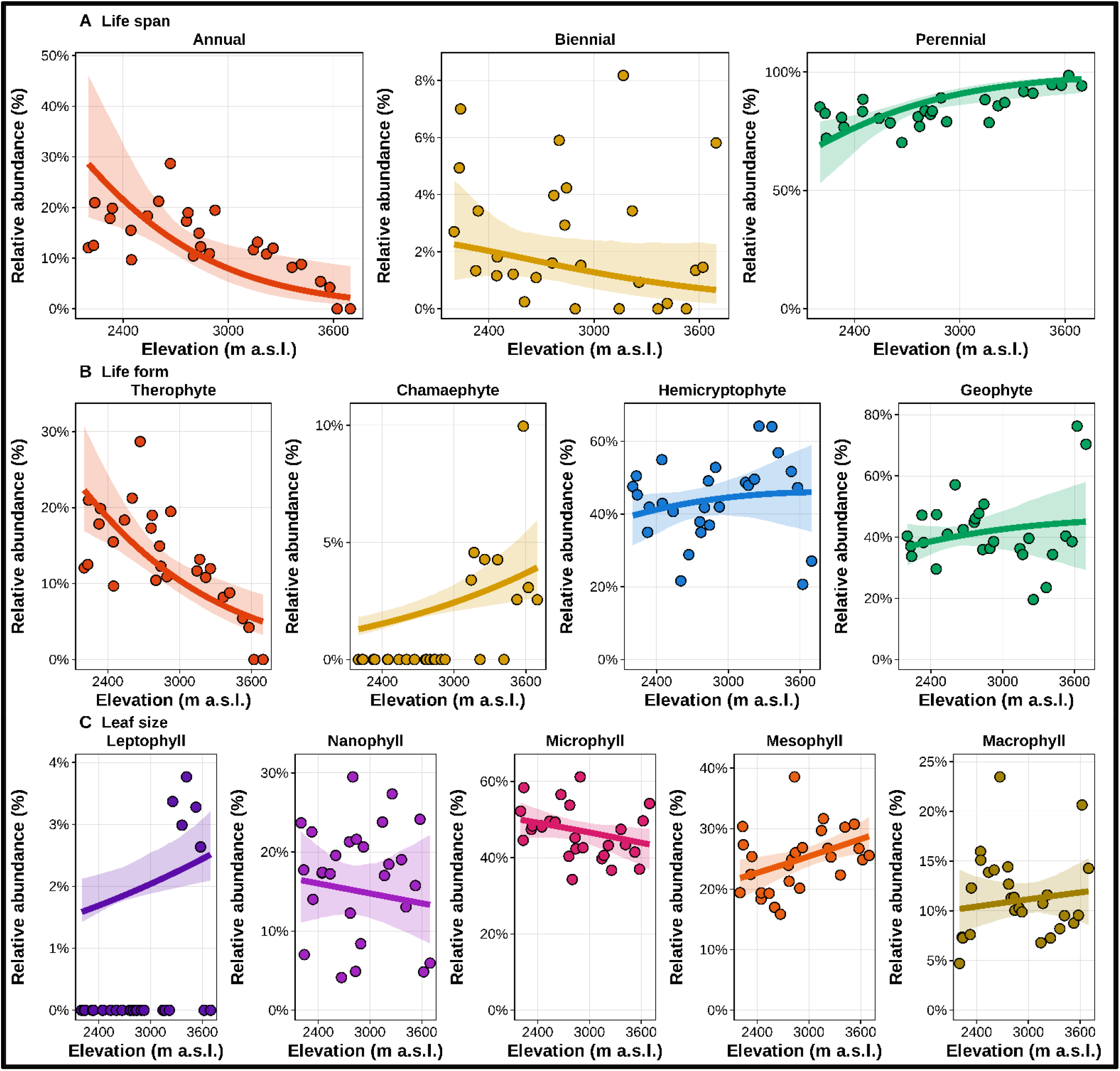
Elevation restructures herbaceous understory functional composition toward perennial and chamaephytic strategies, with a concurrent increase in leptophyll representation. **(A)** Life-span composition shifts from annual toward perennial dominance with increasing elevation. Annual relative abundance declines significantly, whereas perennial relative abundance increases significantly; biennial species show no significant linear elevational response. **(B)** Growth form composition shows the strongest restructuring across the gradient. Therophyte relative abundance declines significantly with elevation, while chamaephyte relative abundance increases significantly. Hemicryptophytes and geophytes show no significant linear responses. **(C)** Leaf-size composition also varies along the elevational gradient, but only leptophyll relative abundance increases significantly with elevation. Nanophyll, microphyll, mesophyll and macrophyll classes show no significant linear responses after multiple-testing correction. Dots represent observed abundance-weighted relative abundances at each study site. Solid lines show fitted elevational responses from class-specific centred log-ratio models, back-transformed and normalized to compositional proportions so that predicted classes within each trait dimension sum to 100%.

Life span and life form show the clearest directional responses (Figure 4A–B). Specifically, perennials increase significantly with elevation (β = 0.999, *R*² = 0.501, *P*adj < 0.001), whereas annuals decline (β = −0.946, *R*² = 0.186, *P*adj = 0.037). Similarly, therophytes show the strongest decline with elevation (β = −0.997, *R*² = 0.597, *P*adj < 0.001), while chamaephytes increase significantly (β = 0.748, *R*² = 0.579, *P*adj < 0.001). In contrast, only leptophylls increase significantly among leaf-size classes (β = 0.235, *R*² = 0.262, *P*adj = 0.032; Table S7). Thus, our findings support H3, with stronger directional shifts in life span and life form than leaf size.

### 3.4 Turnover-driven community differentiation

Our results show that species turnover dominates β-diversity among the 27 study sites (Figure 5A–C; Table S8). Consistent with H4, community differentiation primarily reflects species replacement rather than nested species loss. Total Sørensen dissimilarity is high (βSOR = 0.898), with turnover (βSIM = 0.880) contributing 97.95% and nestedness (βSNE = 0.018) only 2.05%. Specifically, pairwise analysis of 351 site combinations reveals greater mean turnover (0.572) than nestedness (0.038), with turnover exceeding nestedness in 99.1% of pairs (Table S9).

**Figure 5.**
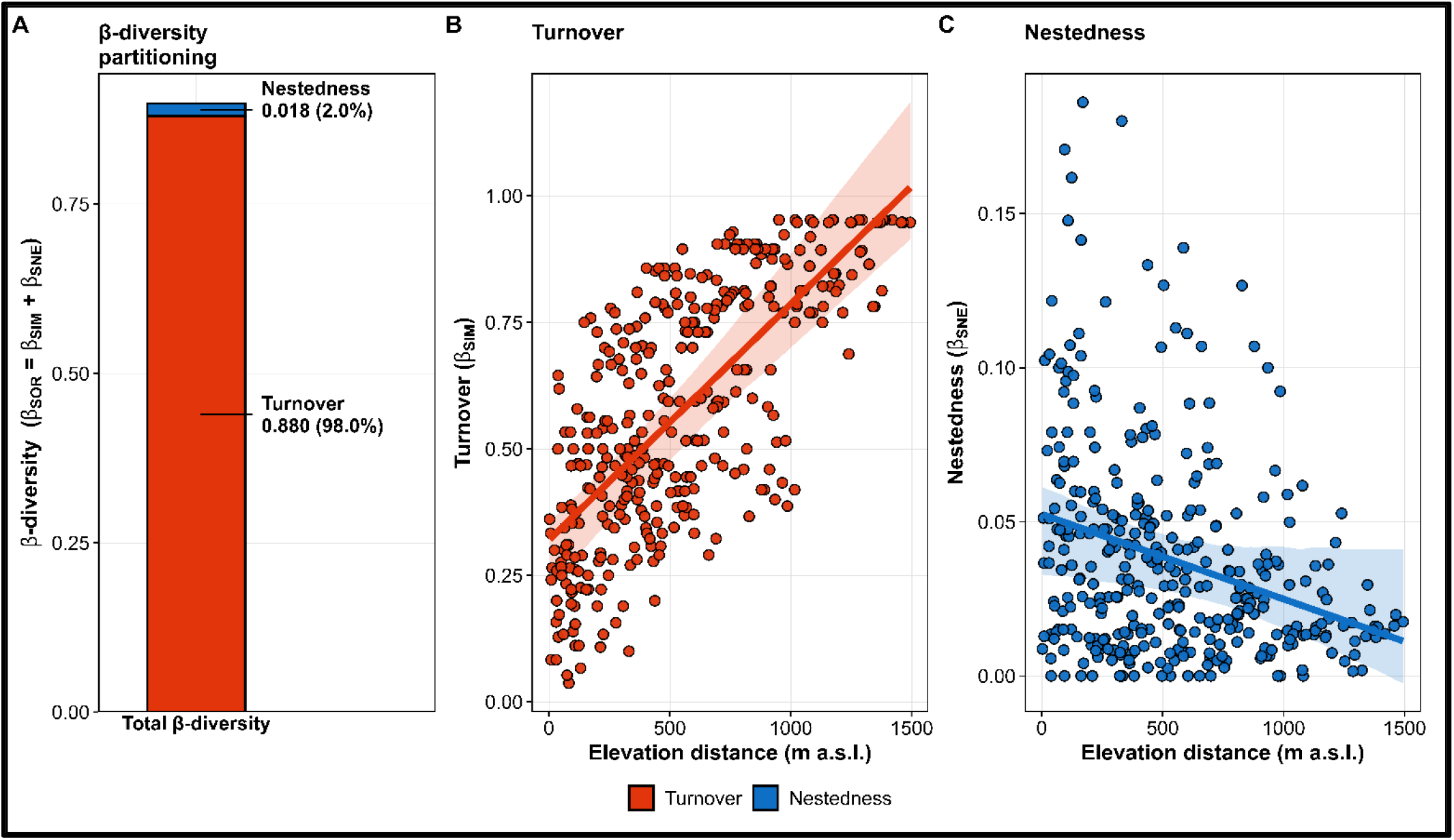
Increasing elevational separation differentiates understory communities primarily through species replacement rather than nested species loss. **(A)** Multi-site Sørensen β-diversity (βSOR = 0.898) is partitioned into turnover and nestedness-resultant components. Species turnover accounts for 97.95% of total β-diversity (βSIM = 0.880), whereas nestedness contributes only 2.05% (βSNE = 0.018), indicating that compositional differentiation is driven primarily by species replacement. **(B)** Pairwise turnover increases with elevational separation across the 351 site pairs, showing that communities farther apart in elevation contain increasingly different sets of species. (**C**) Pairwise nestedness decreases with elevational separation, indicating that compositional differences across larger elevational distances do not arise from increasingly nested species assemblages.

We further found that turnover increases significantly with elevational separation (Figure 5; Table S10). Specifically, pairwise turnover shows a positive relationship with elevational distance (*r* = 0.715, *P* < 0.001), increasing by 0.467 per 1,000 m (95% CI = 0.373–0.560). In contrast, nestedness decreases (*r* = −0.300, *P* < 0.001), declining by approximately 0.028 per 1,000 m (95% CI = −0.039 to −0.016). Thus, our findings support H4, identifying species replacement as the dominant component of community differentiation along elevation.

## 4 Discussion

Mountains support a disproportionate share of terrestrial biodiversity, partly because of their complex terrain and climatic heterogeneity (Antonelli et al. 2018; Rahbek et al. 2019; Perrigo et al. 2020; Manzoor et al. 2026). As climate warming alters mountain vegetation, understanding how elevation structures plant communities is increasingly important (Peters et al. 2016; Rumpf et al. 2018; Steinbauer et al. 2018; Vitasse et al. 2021). Here, we examine 107 herbaceous understory species across 27 sites spanning 2,202–3,696 m above sea level (a.s.l.) in the Kashmir Himalaya. We test hypotheses concerning community differentiation, declining α-diversity, functional shifts, and turnover-driven β-diversity. To do so, we integrate community composition, α-diversity, functional composition, and β-diversity along the same gradient. Our results reveal increasing compositional differentiation and declining local diversity toward higher elevations. Furthermore, perennial and chamaephytic groups increase in representation, whereas annual and therophytic groups decline. By contrast, only leptophylls show a significant positive elevational relationship among leaf-size classes. Notably, species turnover accounts for 97.95% of total Sørensen β-diversity, indicating that differentiation primarily reflects species replacement rather than nestedness. Our findings agree with previous studies reporting diversity declines along elevational gradients (Zhang et al. 2016; Bañares-de-Dios et al. 2024; Irungbam et al. 2025). Likewise, our results align with Wani et al. (2023), who identified elevation as the strongest predictor of community composition and turnover as the dominant component of β-diversity.

Elevation is strongly associated with compositional differentiation among mountain understory communities (Sangry et al. 2024). Our results show that temperate, subalpine, and alpine communities become increasingly distinct with elevational separation, supporting our hypothesis. Multivariate analyses further indicate differences in composition rather than within-group dispersion (Anderson & Walsh, 2013). Our findings agree with Wani et al. (2023), who identified altitude as the strongest predictor of community composition, and Rahman et al. (2023), who reported greater turnover than nestedness along elevation. The observed differentiation may reflect environmental filtering, as reported for woody communities (Bañares-de-Dios et al. 2024). However, elevation encompasses temperature, moisture, and growing-season length (Körner, 2007). Without direct measurements, we cannot distinguish environmental filtering from other assembly processes, including biotic interactions (Kraft et al. 2015; Cadotte & Tucker, 2017). Related evidence comes from He et al. (2023), who found stronger associations of turnover with climatic and topographic differences than spatial distance. More broadly, the observed differentiation aligns with distance-decay patterns (Nekola & White, 1999; Graco-Roza et al. 2022), including increasing turnover with elevational distance (Fontana et al. 2020).

Elevational gradients shape local plant diversity, although these relationships vary across mountain ecosystems (Rahbek, 1995; Ahmad et al. 2025). Our results show significant declines in understory species richness and Shannon diversity with increasing elevation. These findings agree with Wani et al. (2022), who reported declining herbaceous richness along elevation. By contrast, Zheng et al. (2022) found increasing herbaceous richness, whereas Han et al. (2023) reported nonlinear richness patterns among understory herbs. Such differences suggest that regional environmental conditions may influence elevational diversity responses. Importantly, our sampled gradient begins at 2,202 m, so a richness peak below this elevation cannot be excluded. Previous studies have documented mid-elevation peaks and shown that diversity patterns depend partly on the elevational range examined (Grytnes & Vetaas, 2002; Rahbek, 2005; Irungbam et al. 2025). Furthermore, elevation encompasses multiple environmental factors, including temperature and moisture (Körner, 2007). We therefore interpret the observed declines as negative relationships within the sampled gradient rather than evidence of a single underlying ecological mechanism.

Plant life span and renewal-bud position reflect distinct strategies for persistence under unfavorable conditions (Raunkiær, 1934; Salguero-Gómez, 2017). Our results reveal significant elevational shifts in life span, growth form, and leaf-size composition, supporting our prediction. Notably, perennials and chamaephytes increase, whereas annuals and therophytes decline, with therophytes showing the strongest negative response. Our findings agree with Chondol et al. (2023), who reported that long-lived species were common in stressful alpine habitats. Similarly, Di Biase et al. (2021) recorded therophytes mainly at lower elevations, while Midolo et al. (2024) found more therophytes in warm, dry regions and chamaephytes in colder environments. These responses reflect contrasting persistence strategies: therophytes survive unfavorable seasons as seeds, whereas chamaephytes retain protected buds near the ground. However, hemicryptophytes and geophytes show no significant elevational responses, contrasting with Das et al. (2020), who reported increases in hemicryptophytes and cryptophytes toward higher elevations. Leaf-size responses are more restricted, with only leptophylls increasing significantly, consistent with small-leaved alpine flora (Shaheen et al. 2011) and evidence linking smaller leaves to harsher conditions (Wright et al. 2017).

Species replacement and nestedness-resultant dissimilarity represent distinct components of community differentiation (Baselga, 2010). Our results show that turnover dominates understory community differentiation, accounting for 97.95% of total Sørensen β-diversity, whereas nestedness contributes only 2.05%. Pairwise analyses further reveal increasing turnover and declining nestedness with elevational separation. Our findings agree with Soininen et al. (2018), who identified turnover as the dominant component of β-diversity across ecosystems. Similar patterns were reported by Fontana et al. (2020), who found dominant turnover in vascular plants, increasing with elevational distance as nestedness declined. This predominance also aligns with elevational plant-community studies (Cordeiro et al. 2023; Jiang et al. 2023; Loiola et al. 2023). Likewise, Wani et al. (2022, 2023) attributed elevational compositional differences primarily to turnover. In contrast, Liang et al. (2025) reported increasing nestedness with elevation. Together with declining local diversity, turnover dominance indicates that higher-elevation communities are not simply species-poor subsets of lower-elevation assemblages but contain distinct species assemblages.

Conservation planning depends on differences in species composition among sites rather than local richness alone (Socolar et al. 2016). Indeed, compositional differences can increase the complementarity of sites selected for conservation (Beier & Albuquerque, 2016). In this context, our results show that species turnover dominates compositional differentiation, suggesting that conservation assessments should represent communities across the elevational gradient rather than focus exclusively on species-rich sites (Baselga, 2010). Fontana et al. (2020) similarly emphasized considering entire spatial settings in mountain conservation, while Cordeiro et al. (2023) identified turnover as the main component of differentiation along an elevational gradient. Nevertheless, our findings derive from a single survey without direct measurements of climate, soil, or biotic interactions. Repeated surveys are therefore important for detecting temporal change, as long-term studies have documented increasing summit richness and upslope shifts in plant distributions (Steinbauer et al. 2018; Vitasse et al. 2021). Accordingly, conservation planning should consider temperate, subalpine, and alpine communities, including high-elevation habitats supporting specialized species (Wambulwa et al. 2021; Jiang et al. 2023).

## Conclusions

Elevation structures understory plant communities through changes in species identity and functional strategies, rather than local richness alone. In the Kashmir Himalaya, higher-elevation herbaceous communities contain distinct species assemblages with greater representation of perennials and chamaephytes, despite declining local diversity. Notably, species turnover dominates compositional differentiation, indicating substantial species replacement across the elevational gradient. However, elevation integrates multiple environmental gradients, and our analyses cannot isolate the ecological mechanisms underlying these patterns. The observed differentiation is particularly relevant as climate warming continues to reshape mountain vegetation. Indeed, Steinbauer et al. (2018) documented accelerating species richness gains on mountain summits, while Rumpf et al. (2018) reported delayed range shifts and greater losses among cold-adapted high-elevation species. Together, these findings highlight the potential vulnerability of distinct high-elevation communities and the importance of baseline surveys for detecting future changes. Our study therefore contributes to a broader understanding of mountain biodiversity patterns (Rahbek et al. 2019b) and provides a foundation for long-term ecological monitoring and conservation across elevational gradients.

## Author contributions

Author contributions: SWG and RSG designed the research. SWG performed the analyses and wrote the manuscript. SWG designed the methodology and conceptualized the study with input from RSG and MM. SWG and MM collected the field data. MA and RSG supervised the study. RSG analytically reviewed, edited and improved the initial draft. SWG led the first draft of the manuscript under the guidance of RSG. MM and MA data curation and visualization. All authors edited the draft for final submission.

## Acknowledgments

We thank the local communities of the Himalayan region for their invaluable assistance during fieldwork. We also acknowledge the Department of Plant Sciences, Quaid-i-Azam University, Islamabad, Pakistan, and the Department of Biology, University of Oxford, United Kingdom, for providing support during initial data collection and access to facilities for manuscript completion. SWG was supported by a Commonwealth Split-Site Scholarship at Oxford, hosted by RSG. RSG was supported by a NERC Pushing the Frontiers grant (NE/X013766/1).

## Data availability statement

The supporting findings of this study are provided in the Supplementary Material accompanying this article.

